# A low-cost DIY device for high resolution, continuous measurement of microbial growth dynamics

**DOI:** 10.1101/407742

**Authors:** Kalesh Sasidharan, Andrea S. Martinez-Vernon, Jing Chen, Tiantian Fu, Orkun S Soyer

**Affiliations:** School of Life Sciences, University of Warwick, Coventry, CV4 7AL, UK; Synthetic Biology Centre for Doctoral Training, University of Warwick, Coventry, CV4 7AL, UK; Warwick Integrative Synthetic Biology Centre (WISB), University of Warwick, Coventry, CV4 7AL, UK

**Keywords:** Microbial growth, automated measurement, optical density, synthetic biology, growth rate fitting, microbial growth model, Arduino, citizen science, anaerobic culturing

## Abstract

High-resolution data on microbial growth dynamics allow characterisation of microbial physiology, as well as optimisation of genetic alterations thereof. Such data are routinely collected using bench-top spectrophotometers or so-called plate readers. These equipments present several drawbacks: (*i*) measurements from different devices cannot be compared directly, (*ii*) proprietary nature of devices makes it difficult for standardisation methods to be developed across devices, and (*iii*) high costs limit access to devices, which can become a bottleneck for researchers, especially for those working with anaerobic organisms or at higher containment level laboratories. These limitations could be lifted, and data reproducibility improved, if the scientific community could adopt standardised, low-cost and open-source devices that can be built in-house. Here, we present such a device, MicrobeMeter, which is a do-it-yourself (DIY), simple, yet robust photometer with continuous data-logging capability. It is built using 3D-printing and open-source Arduino platform, combined with purpose-built electronic circuits. We show that MicrobeMeter displays linear relation between culture density and turbidity measurement for microbes from different phylogenetic domains. In addition, culture density estimated from MicrobeMeter measurements produced less variance compared against three commercial bench-top spectrophotometers, indicating that its measurements are less affected by the differences in cell types. We show the utility of MicrobeMeter, as a programmable wireless continuous measurement device, by collecting long-term growth dynamics up to 458 hours from aerobic and anaerobic cultures. We provide a full open-source description of MicrobeMeter and its implementation for faster adaptation and future development by the scientific community. The blueprints of the device, as well as ready-to-assemble kit versions are also made available through www.humanetechnologies.co.uk.

## INTRODUCTION

One of the key measurements in microbiology, and the associated fields of systems and synthetic biology, is the growth rate of individual microbial species. This measurement provides qualitative confirmation on the types of substrates and conditions a microbe can grow in, and can be used to infer quantitative growth rates under such conditions(1, 2). The resulting information is essential for building predictive models of microbial growth and understanding the impact of genetic or environmental alterations, as well as the optimisation of media conditions or genetic modifications. Therefore, measurement of microbial growth dynamics is utilised in all microbial, synthetic and molecular biology laboratories, and within the biotechnological and biomedical industries.

For the actual measurement of growth dynamics, scientists generally use bench-top spectrophotometers or so-called plate readers (which implement similar optics as spectrophotometers). These devices are used in measuring cell cultures’ absorption, typically at a wavelength of 600nm, and the resulting values are reported as culture ‘turbidity’ or ‘optical density (OD)’. Technically, however, spectrophotometers are designed to measure light absorption of small molecules, where the measured absorption value relates linearly to the concentration of the molecule and the path length of the light through the container, as captured by the Beer-Lambert law (3). It is well-known that this law does not fully apply to large colloids suspended in liquid, such as microbial cells, as these larger particles can display both absorption and scattering of light (4–6). Thus, reported turbidity values are not absorption values, but rather a combination of absorption and scattering of light through the cell culture (4, 5, 7). The ratio between scattering and absorption of cell cultures can be affected by various factors such as cell size, type of cell membrane, growth medium, cell density, path-length, and the presence of pigments (5, 7–9). The scattering effects can also interact with the specific optical design of a given spectrophotometer, producing differential effects on the resulting measurements between spectrophotometers with different optical designs (4, 5, 7, 10, 11). As expected from these considerations, turbidity measurements of the same cell culture samples using different spectrophotometers are found to have significant variability (sometimes more than 30%) (5, 10, 11). The relation between the turbidity and cell concentration, while linear, is also found to vary from device to device (5–7). These results show that both reported turbidity values and the turbidity-based calculation of microbial growth rates are device specific.

This presents a significant challenge for reproducibility. In practical terms, any experiment requiring exact modulation of microbial culture concentration cannot be accurately reproduced, unless one uses the same spectrophotometer that is used to generate the original concentration information. One possible solution to this challenge is to ‘calibrate’ all spectrophotometers against a chosen standard solution. This standard solution could be used for calculating the ratio between the turbidity measurements obtained using two different spectrophotometers, and then applying that ratio as a conversion factor between the devices. In practice, however, such conversion factors are found to be highly specific to individual samples that are used for obtaining them. For example, the conversion factor obtained from a standardised solution, such as the McFarland standard (12), is not applicable in the case of corrections required to be applied when measuring cell cultures (5, 7, 10, 11). This creates a practical difficulty in establishing conversion factors between different devices, where different conversion factors would need to be calculated among all devices and on different samples.

An alternative approach to achieving standardisation and reproducibility of microbial growth measurements would be for the scientific community to agree on the use of a single photometric device for this measurement. This would allow collection of data under the same optical design, and thereby reducing the reproducibility problems that are associated with device-to-device variability. For it to be adaptable by as many research groups as possible, such a ‘universal’ turbidity measurement device should ideally be simple in construction, low-cost, open source, and robust. These properties are not met by current bench-top spectrophotometers, and the more high-throughput plate readers, as these are relatively high cost devices and their manufacturing and optical design details are proprietary. In addition, the optics of these devices are optimised for purposes other than just monitoring cell culture dynamics, increasing the design complexity (and possibly the effects of scattering). A simple, low-cost and open-source device designed solely for cell growth measurement could thus allow increased reproducibility of experiments relating to microbial growth and also open-up scientific analyses to larger groups of people through reducing the cost of required scientific instrumentation, while supporting the philosophy of do-it-yourself (DIY) science (13, 14).

Here, we develop such a DIY device for automated monitoring of microbial growth dynamics under both aerobic and anaerobic conditions. This device, called MicrobeMeter, is developed using open-source Arduino and 3D printing technology and software, resulting in an estimated single-unit production cost under £150. These specifications and the modular nature of MicrobeMeter’s design are expected to allow further development and optimisation of the device by the research community, as well as its possible adaption as a ‘universal’ turbidity measurement device. We demonstrate the feasibility of MicrobeMeter to capture microbial growth dynamics by measuring the turbidity of the serial dilutions of three different organisms: a facultative aerobe (*Escherichia coli*), a heterotroph (*Shewanella oneidensis*), and a yeast (*Schizosaccharomyces pombe*). MicrobeMeter produced linear measurements (R^2^ ≥ 0.99) for the dilution series of all these cultures, and the slopes of the measurements were ~92% close to the calculated theoretical slopes. Compared against commercial bench-top spectrophotometers, MicrobeMeter measurements produced the least variance in slope, indicating that the turbidity measurement of the device is less affected by the differences in cell types. In addition, the device-to-device variability among MicrobeMeter replicates was very low, indicating robustness towards manufacturing errors. We also demonstrate the use of MicrobeMeter as a continuous measurement device under aerobic and anaerobic conditions, by collecting long-term growth dynamics of *E. coli* (under aerobic conditions), and two strictly anaerobic microbes: a sulfate-reducing bacterium (*Desulfovibrio vulgaris*) and a methanogenic archaeon (*Methanosarcina barkeri*). Taken together, these results show that MicrobeMeter could be used as a reliable, simple-to-construct and cost-effective photometer for the turbidity measurements of cell cultures and derivation of kinetic modelling parameters relating to growth dynamics. All information regarding the manufacturing of MicrobeMeter (3D models of device casing, electronic circuits and component list, and software) is made available for personal and academic non-commercial research use through this publication and from a dedicated website (see *Device Availability* section), allowing this simple photometer to be reproduced and adapted by the scientific community.

## RESULTS

As the use of complex optics contributes significantly to the disparities between commercial spectrophotometers (4, 5, 10, 11), an ideal universal measurement device should be devoid of complex optics to increase its reproducibility and cost-effectiveness. It is important to note that although spectrophotometers are capable of taking measurements at different wavelengths of light, turbidity measurements of cell cultures are widely conducted using 600nm wavelength (hence, most reports refer to the resulting turbidity measurements as OD_600_).

### MicrobeMeter has a simple design using low-cost components

Using the above fact allowed us to simplify the optics in turbidity measurement by using a single 601nm light emitting diode (LED), a suitable light-sensor (a silicon photodiode) and two simple apertures as the sole optical components in MicrobeMeter (Figure 1A, and see *Methods*). These components are integrated in a 3D-printed casing, which also acts as a holder for the culture vessel (Figure 1B). The photodiode voltage is measured using a purpose-built, Arduino integrated electronic circuit board that acts as an analogue to digital converter (ADC) with high sensitivity (see below and also *Methods*). Using the Bluetooth compatibility of the Arduino, the acquired data are sent wirelessly to a computer, where they can be further analysed.

The current design is built around a standard test tube as the culture vessel, rather than a quartz cuvette (note that the use of quartz cuvettes is primarily for achieving better optics in absorption measurements, which is not the primary measurement in cell cultures as discussed above). The use of a test tube allows for larger culture volumes to be used, and for continuous measurements to be obtained directly during growth experiments (minimum measurement interval is set at 6s; see *Methods*). This approach also easily accommodates anaerobic experiments using sealed test tubes with the dimensions of a standard test tube, such as Hungate tubes, which we used as the standard for our current design (see *Methods*). The tube holder is designed to shield the tube from external light to minimise the interference to the measurement. Two identical apertures were created at the light source and sensor sides. The aperture diameter was set to 2mm, which is possible to achieve with even low-cost 3D-printers. The design of MicrobeMeter consists of ports for four-tubes and accommodates all electronic components including a battery, and it is held in a compact container for further protection (Figure 1B; see also *Methods*).

**Figure 1:**
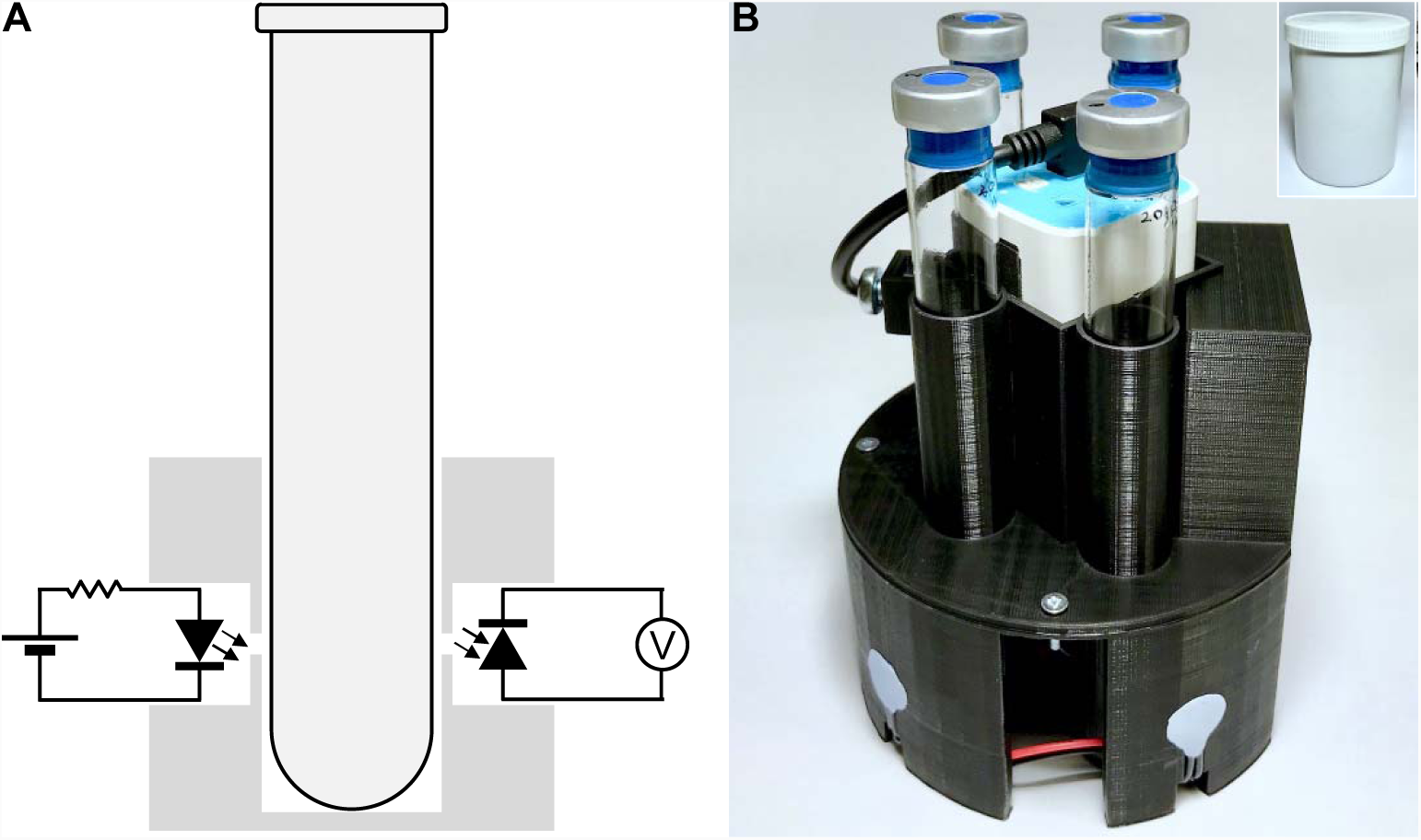
A: Schematic showing the cross section of a simple photometer. The dark grey region shows the plastic body of the device that holds a LED (left), a photodiode (right), and a test tube (middle). The LED is connected to a battery via a resistor. The photodiode is connected to a volt meter for measuring the voltage generated by the photodiode upon the incident of the light from the LED through the apertures. **B: The design of MicrobeMeter.** The picture shows (in black) the 3D-printed casing of MicrobeMeter, holding the measurement tubes (i.e., samples and blank), battery, and Arduino compatible electronic circuit board. The casing and electronic circuit designs are available for personal and academic non-commercial research use through a dedicated website (see *Device Availability*). Insert at top-right corner shows MicrobeMeter placed in a white plastic jar for further protection.

### MicrobeMeter is designed for robust turbidity measurements with high signal-to-noise ratio

To achieve high signal-to-noise ratio and low variability in MicrobeMeter measurements, we implemented: (*i*) a state-of-the-art amplifier circuit on the Arduino-compatible electronic circuit board, (*ii*) an averaging function into the data acquisition program, and (*iii*) an optimised measurement routine that stabilises the analogue to digital converter (ADC) output of the Arduino (see *Methods* for details). The electronic circuit board design implements a chopper-stabilised operational amplifier (15) connected as a trans-impedance amplifier with common-mode rejection and differential input configuration. This configuration offers sensitive measurements that are highly immune to external noises arising from electrostatic coupling (16). This amplifier is set to have a gain of 10,200,000 and an output range of 0-5V.

Given this amplifier and optimised measurement routine, we determined the linearity of response in a technical quality control experiment. Using the MicrobeMeter, we took independent measurements on each port, while reducing the LED light intensity from maximum to minimum using Arduino’s built-in pulse-width modulation (31372.55Hz) feature. Performed with Hungate tubes containing 10mL of distilled water, these controlled measurements mimicked growth of a biological cell culture (and the resulting reduction in transmittance). Raw light intensity measurements were converted to turbidity values using –log(*I*_*t*_/*I*_*0*_), where *I*_*t*_ is the light intensity at time *t*, and *I*_*0*_ is the light intensity at time 0. As shown in Figure 2, measurements collected on each port demonstrated near-perfect linearity, while variability between ports was low (see *Supplementary File 1* for raw values). The raw light intensity values measured by each port was always within measurement range of the device (see *Methods*). To allow users to check their MicrobeMeter unit, this technical quality control experiment is integrated in the data acquisition program (see *Methods*).

**Figure 2:**
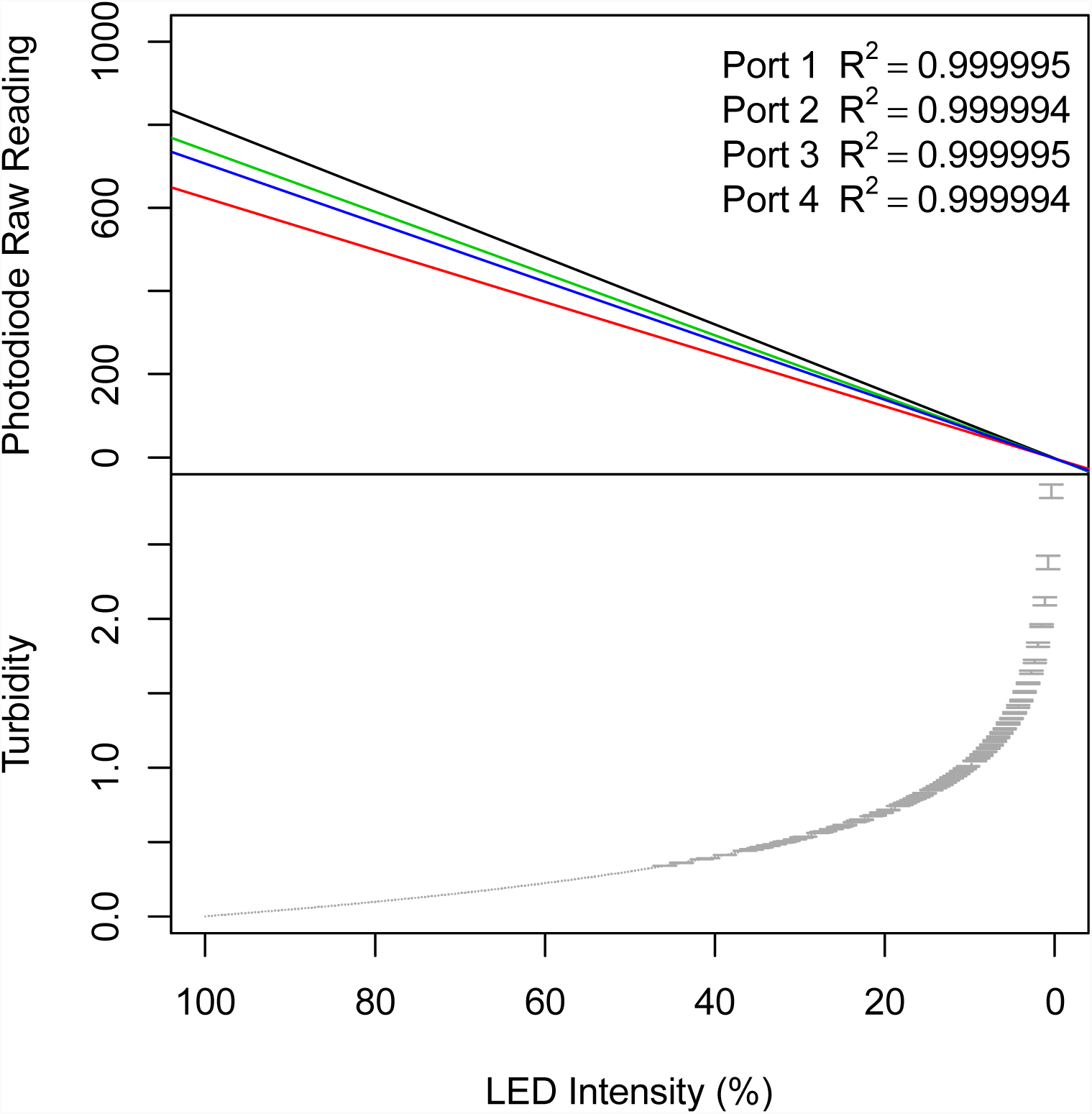
Linearity of MicrobeMeter response to light. The linearity of the response of the four ports of MicrobeMeter was determined by lowering the light intensity from its maximum to minimum by adjusting the duty cycle of the signal that drives the LED (x-axes), while recording the photodiode raw readings (y-axis of top panel). The coefficient of determination (R^2^) is shown in the legend. The raw photodiode readings from each port (top panel) were converted to turbidity values (see main text), the average (black dots) and the standard deviation (grey bars) of which are shown in the bottom panel.

### MicrobeMeter based estimation of cell culture density from turbidity is comparable to, or better than, bench-top spectrophotometers

Having shown the ability of MicrobeMeter to achieve high sensitivity with simulated turbidity measurements of blank samples, we next tested the linearity of response using biological samples. To this end, we used cell cultures of three different organisms that are expected to have different scattering properties: *E. coli, S. oneidensis* and *S. pombe* (see *Methods*). For each organism, two-fold serial dilutions were prepared by harvesting the cells via centrifugation and re-suspending in standard phosphate-buffered saline (PBS). Each cell dilution series was measured using MicrobeMeter with three tubes containing the same culture sample and one tube containing PBS only (i.e., “blank” measurement) and used for temperature correction (see *Methods*). The same serial dilutions were also measured using three different bench-top spectrophotometers (see *Methods*). For all tested cell culture dilutions, the resulting turbidity measurements were linear both for MicrobeMeter and for the commercial spectrophotometers (R^2^ ≥ 0.99) (Figure 3A). As expected, all devices resulted in different actual values from each other (Figure 3A), confirming that the turbidity measurements cannot be compared between devices (5, 10, 11).

The average of the slopes of the three serial dilutions was compared against the calculated slope of the serial dilution, which is expected to be *ln*(2) given the two-fold dilutions performed. The averages of the slopes produced by the three ports of MicrobeMeter displayed lower variability compared to slopes obtained from commercial spectrophotometers (Figure 3B). These results show that the turbidity measurements obtained using MicrobeMeter allows estimating cell density as accurately as commercial spectrophotometers and provides less variability across estimates. The latter finding suggests that MicrobeMeter is less sensitive to the differences in scattering caused by the different tested cell types in comparison to the spectrophotometers tested.

**Figure 3:**
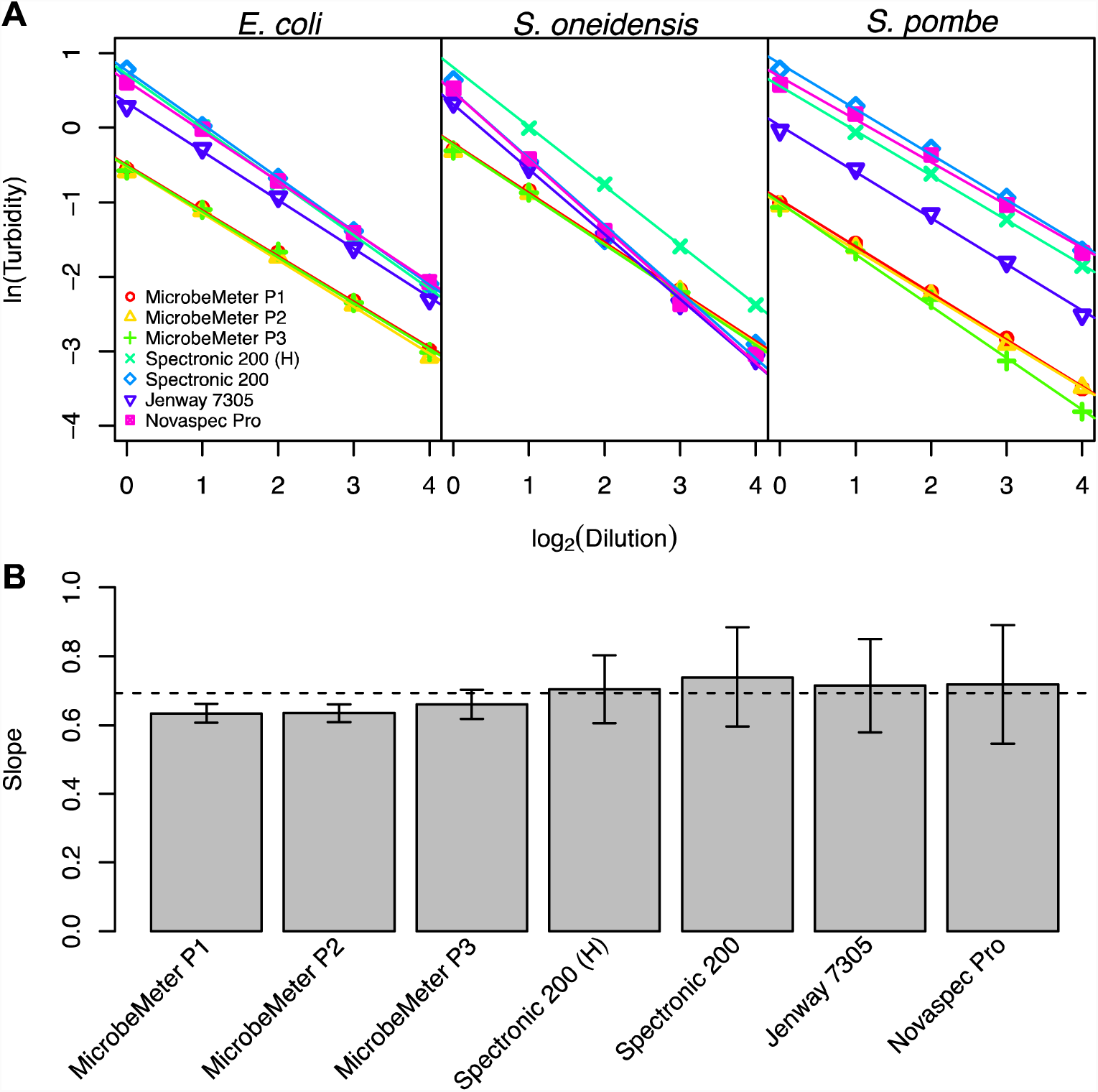
A: Regression analysis of the turbidity measurements taken from the serial dilutions of three different cell cultures (plot titles). Measurements are obtained using the MicrobeMeter (P1, P2 and P3 indicate the three ports of MicrobeMeter) and three commercial spectrophotometers. The x-axes show culture dilution levels, while the y-axis shows the natural logarithm of the blank subtracted and path-length corrected turbidity values. Regression line was calculated for each dataset as shown. The coefficient of determinations (R^2^) of all datasets were greater than 0.985 (not shown in the figure). The Spectronic 200 was used for obtaining measurements from samples kept both in cuvettes and Hungate tubes (H). **B: The slopes of the turbidity measurements of the serial dilutions.** Each bar shows the average and standard deviation (error bar) of the slopes of three different cell cultures shown in panel A. The dotted line indicates the calculated slope of the serial dilutions (as described in the main text).

### MicrobeMeter can be used for long-term continuous monitoring of growth dynamics under aerobic and anaerobic conditions

Following on from the measurements of cell culture serial dilutions, we next tested the ability of MicrobeMeter to be used in continuous monitoring of cell culture growth. This overcomes the need for manual sampling and allows automated acquisition of high-resolution growth dynamics. Furthermore, the wireless capability of MicrobeMeter allows it to be placed in an incubator, clean bench, containment or anaerobic chamber without needing to run wires for power supply and data acquisition.

To test long-term growth dynamics measurements, we used MicrobeMeter with open or sealed Hungate tubes. We then collected high-resolution turbidity data under aerobic conditions for the facultative organism *E. coli* (Figure 4), and under anaerobic conditions for the strictly anaerobic organisms *D. vulgaris* and *M. barkeri* (Figure 5). Measurement periods for these organisms were approximately 30.5, 75, and 458 hours, respectively, allowing us to collect unprecedented growth dynamics data. For the case of *E. coli*, we compared this high-resolution data with measurements of the technical replicates of the same cultures, obtained using a commercial bench-top spectrophotometer (see *Methods*). Applying an appropriate conversion factor (see *Methods*), the measurements using the commercial spectrophotometer and MicrobeMeter showed near perfect overlap for the *E. coli* data (Figure 4). This demonstrates the reliability and potential application of MicrobeMeter as a continuous turbidity measurement unit to be used in the laboratory setting, under both aerobic and anaerobic conditions.

**Figure 5:**
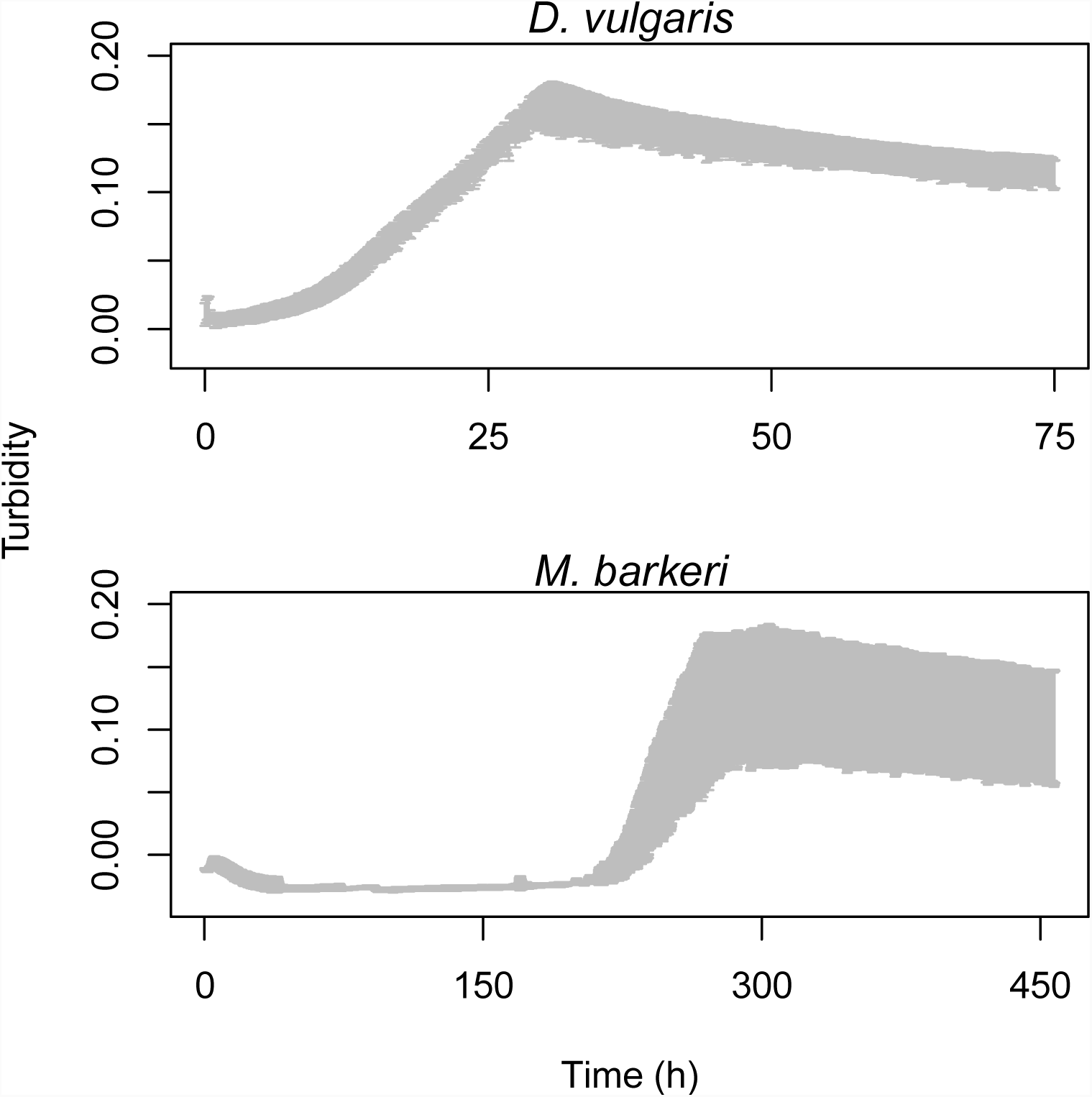
Continuous turbidity measurements of *D. vulgaris* and *M. barkeri* cultures using MicrobeMeter. Long-term turbidity measurements were taken using three technical replicates of *D. vulgaris* and *M. barkeri* cultures. The black dots and grey bars show the average and standard deviation of measured turbidity values, respectively. The x-axes show the time in hours. All turbidity values are blank subtracted and path-length corrected. Smoothing, using a moving average with window size of ten, was performed on the measurements of *M. barkeri* cultures. The MicrobeMeter measurement frequency of *D. vulgaris* and *M. barkeri* cultures was five minutes.

**Figure 4:**
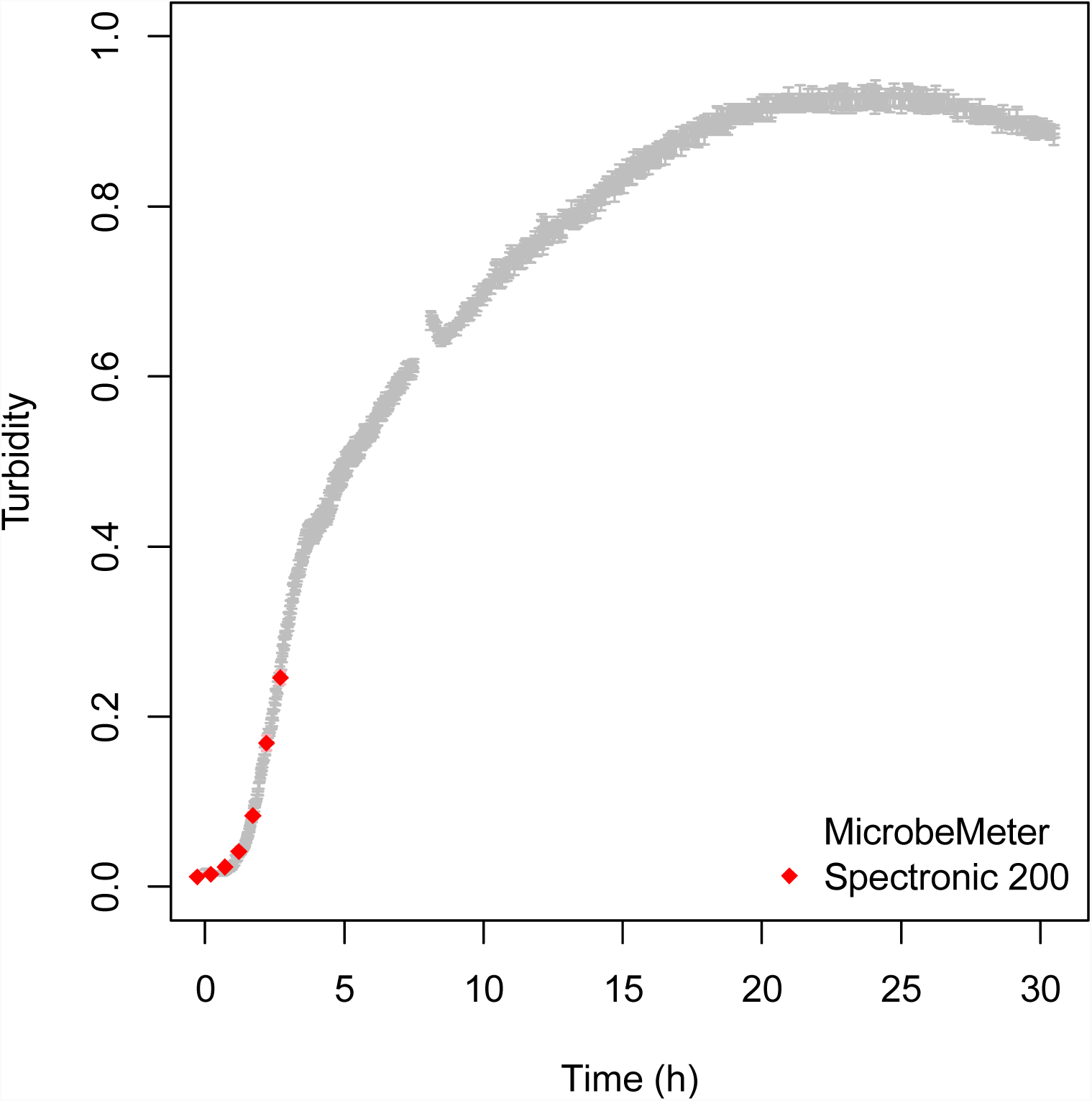
Continuous turbidity measurements of *E. coli* cultures using MicrobeMeter. Long term turbidity measurements were taken using three technical replicates of *E. coli* culture. The black dots and grey bars show the average and standard deviation of measured turbidity values, respectively. The x-axis shows the time in hours. These data were compared against manually collected data on an additional set of three technical replicates of *E. coli* culture, obtained using a commercial spectrophotometer (Spectronic 200; average: red rhombus; standard deviation: light red bars). Note that the Spectronic 200 measurements were multiplied using a conversion factor (as described in the main text) to fit with the MicrobeMeter measurements. All turbidity values are blank subtracted and path-length corrected. The MicrobeMeter measurement frequency was one minute. A gap was introduced in the measurements, at about 8h, when the cultures were removed for visual inspection.

## DISCUSSION

Here, we present a low-cost DIY photometer called MicrobeMeter. It consists of LEDs, photodiodes, and a 3D-printed casing that can hold four test tubes, a battery and a purpose-built Arduino-compatible electronic circuit board for signal amplification, noise cancellation, and data logging. We show that MicrobeMeter can perform linear measurements on a range of serial dilutions of microbial cells cultures, and that these measurements are as good as, or better than those obtained from common commercial bench-top spectrophotometers. We demonstrate the utility of MicrobeMeter by collecting long-term continuous measurements of microbial cultures both under aerobic and anaerobic conditions. These measurements can be taken in a shaking incubator or anaerobic cabinet, given the small footprint of MicrobeMeter. The fully disclosed design and construction information of MicrobeMeter bring these features to any researcher or citizen scientist at a cost of around £150, presenting MicrobeMeter as a potentially adaptable device for turbidity measurements by the scientific community.

Besides its cost advantage, the open-source and simple nature of the presented device can have significant impact in terms of data reproducibility. Despite the wide spread use of turbidity measurements of microbial cultures, this measurement remains as one that cannot be directly compared between devices (Figure 3) (4, 5, 10, 11). Given its low cost and simplicity, MicrobeMeter, or similar devices, could become ‘universal’ in the sense that they can be employed across many research labs and allow the acquisition of comparable turbidity measurements. To this end, we note that several other DIY-style photometers have also been described. For example, a simple photometer design is implemented on a flow cell for monitoring the turbidity of continuous and/or fed-batch microbial cultures (5). Another simple design was developed for the measurement of flask and/or test tube grown cell cultures (11). The latter device offers also UV-Vis spectroscopy capabilities; however, it makes use of several proprietary components.

The presented device, MicrobeMeter, is designed to be cost effective, open-source and customisable. These features allow it to be adapted by low-income, as well as DIY biology research groups. Both communities are indeed in need of reliable and accurate measurement tools at a low cost and with an open source nature (13, 14). The latter is an important feature, as users should be able to modify such devices in a variety of ways to fit their needs. MicrobeMeter provides such adaptability and flexibility as its electronics are built on the popular and cost-effective Arduino platform, and its physical design can be readily modified in different ways. The 3D casing, for example, can be replaced by simpler and cheaper wood or cardboard design, or its tube-holders can be adapted for different cell culture vessels or use it as flow cells. Similarly, the electronics can be integrated with a display panel to use it as a traditional photometer, while measurement spectrum can be extended by inclusion of LEDs with different wavelengths.

The open-source nature of DIY devices such as MicrobeMeter can also encourage innovation in education and in scientific research. In particular, it is not uncommon that measurement devices developed for a particular measurement with a particular experiment in mind can outperform multi-purpose devices or open up new measurement areas (e.g. see (17)). In the case of MicrobeMeter, we hope that the wireless data communication and programmable continuous measurement features will allow for collection of high-resolution microbial growth data, which can be shared at online databases and contribute to the development of better mathematical models of microbial physiology. Such resources, as well as the DIY nature of the data collection devices, could help train interdisciplinary scientists who have a better grasp of measurement techniques, the resulting data, and their possible uses across scientific borders.

## METHODS

### Design of MicrobeMeter casing

The MicrobeMeter casing was designed using OpenSCAD version 2015.03-3. It was then manufactured using a 3D-printer (Ultimaker 2/2^+^, 0.4mm nozzle; Ultimaker B.V., Geldermalsen, The Netherlands) and polylactic acid filaments (2.85mm diameter; Ultimaker B.V., Geldermalsen, The Netherlands). The 3D-printer was operated as per manufacturer’s instructions and using the following parameters: 0.1mm layer height, 20% infill and with support structure enabled. The photodiode and LED were placed in tunnel-shaped holding bays, with identical apertures at both ends (2mm in diameter and 3mm in length), while the culture tubes were placed in cylindrical holders with an inner diameter of 18.5mm. Thus, the total light path length is 24.5mm. The centre of apertures was placed 15.9mm above from the bottom of the culture tubes (Figure 1A). The tube holder size was optimised to fit a standard Hungate tube (CLS-4209-10, anaerobic culture tubes, Chemglass Life Sciences, New Jersey, USA) with an outer diameter 18mm, which is also the diameter of several other regular test tubes. The 3D design information of MicrobeMeter are provided for personal and academic non-commercial research use through a dedicated website (see *Device Availability* section). The casing design specifications can be changed if needed by the user (e.g. to fit different growth vessels and tubes).

### Optical and electronic components

The complete MicrobeMeter parts list, along with the electronic circuit diagram is provided for personal and academic non-commercial research use through a dedicated website (see *Device Availability* section). A LED with specific wavelength of 601nm (L-53SED, Kingbright, New Taipei, Taiwan) was used as the light source and a silicon photodiode (BPW21R, Vishay, Selb, Germany) was used as the light sensor. Both the LEDs and the photodiodes were controlled (i.e., illumination and measurement, respectively) using an electronic circuit that consists of regulators for LEDs, amplifiers for photodiodes, a temperature sensor, and a Bluetooth data communication module. The electronic circuit is designed as a “shield” to work with Arduino Mega microcontroller (Arduino Mega 2560 Rev3). Each MicrobeMeter unit was powered by a dedicated battery (TL-PB10400, 10400 mAh Power Bank, TP-Link, Reading, UK) which, along with the device, can be packed in a plastic container (PJSCPP1.75LW_BUNDLE; height: 165.3mm; diameter: 126.2mm) (see Figure 1B). In multiple tests, the maximum runtime of MicrobeMeter on a fully charged battery was ~100 hours (measurement frequency was set to 5 minutes). A step-by-step instruction of the MicrobeMeter assembly is provided for personal and academic non-commercial research use through a dedicated website (see *Device Availability* section).

### Data acquisition and processing

MicrobeMeter can acquire data in an autonomous fashion and send it to a computer. This is achieved by the above said Bluetooth enabled Arduino Mega shield and two dedicated programs, written in a server-client architecture. The server component is written using Arduino Software version 1.8.3 and is interfaced with a client component that is written using Perl version 5.18.2. The client program is currently optimised to run on macOS version 10.13.1 and Windows version 8 & 10, and can be readily adapted to other operating systems. The server program runs on the Arduino microcontroller and allows wireless connection to the client program to send the measurements made. Source code for both programs are provided for personal and academic non-commercial research use through a dedicated website (see *Device Availability* section).

The Bluetooth connection between MicrobeMeter and a computer can be established using standard methods as per its operating system (tested using MacBook Pro and MacBook Air with macOS version 10.12 & 10.13, and desktop and laptop computers with Windows version 8 & 10). The default Bluetooth device name of MicrobeMeter is “HC-06” and the password is “1234”. The Bluetooth connection needs to be configured only once, when connecting with the same computer. Once a Bluetooth connection is configured, the client program can be executed on the computer to start individual or long-term measurements with MicrobeMeter. When the program initiates, it proceeds first with setting up user-defined specifics, such as experiment name and time between measurements, then initiates the collection of ‘blank’ and sample measurements. The measurement is taken from one port at a time and the minimum interval between each set of measurements (i.e., from four ports) is set at 6s. This setting can be altered by the user, if desired. The measurements will continue until the user terminates the client script or turns off MicrobeMeter. In case of Bluetooth connection failure or interruption at the computer side, the client program will seek and re-establish connection with MicrobeMeter when the computer is back online. At the beginning of an experiment, the client program creates two files that store the connection status and the measurements. The latter file contains the version of the device, experiment name, measurement time, temperature, and the measurement data. While users can develop appropriate post-analysis approaches for this data, an R script for basic analysis and plots is provided for personal and academic non-commercial research use through a dedicated website (see *Device Availability* section).

### Measurement range and conversion factor

The maximum light intensity that the device can measure is effectively its “measurement saturation point”. Given its use of the Arduino platform, this point is theoretically at 5V (corresponding to an integer reading of 1023) for MicrobeMeter. The design is optimised, so to have a maximum of light reading below 1023 (on the Arduino platform). To ensure this, the Arduino shield circuit is implemented in a way so that when using MicrobeMeter with a Hungate tube containing 10mL distilled water, the maximum intensity light setting produces an integer reading between 600-1000. This allows the device maxima to be below the saturation point of 1023, while still allowing a minimum integer reading range of 1-600 (or up to 1-1000), which corresponds to more than 2.78 turbidity units (e.g., -log(1/600)). In addition, this setting allowed us to account for the observed lensing effects from water-filled Hungate tubes, which can cause approximately 1.6-fold increase in light intensity. As shown in Figures 3 to 5, the resulting measurement range is sufficient to cover detection of microbial cultures’ growth from very early stages well into stationary phase, and the measurements are linear in this range.

For two photometers producing two different measurements for the same sample, a simple conversion factor can be calculated by taking the ratio of the two measurements (10). However, as photometers are differently sensitive to the optical properties of the samples, the conversion factor can vary as the sample changes its properties, for example as changes occur to the cells and medium during a growth experiment (7). Such a normalisation based on a conversion factor was used in Figure 4, where the average of conversion factors over all time points was used for the normalisation.

### Further specifications of MicrobeMeter

The additional aspects of the device are as follows; ***Response time:*** The response time of MicrobeMeter (i.e., the time required for the device measurement to equilibrate after a full rise or fall of signal) was determined by exposing the photodiode for 10s each to the maximum and minimum light intensity of the LED, allowed by the MicrobeMeter circuit (i.e., a cycle). These measurements were done using a Hungate tube containing 10mL distilled water, and with the cycle being repeated for six times at room temperature (see *Supplementary File 2* for the raw data). The response times are determined by taking the average of the device output during the second half of each signal as the target value, and identifying the time when that target value is reached after a maximum/minimum signal is originally initiated (a moving average with window size ten was used to smooth the signal). The longest response times recorded among any of all tested ports were 57ms and 31ms, respectively, for raising (i.e., minimum to maximum) and falling (i.e., maximum to minimum) signals. These results suggest that 88ms plus the time for measurement should be the minimum time gap between two independent measurements on MicrobeMeter. Currently, the default value for this gap is set to 0.9s, well-above this suggested minima. This setting can be altered by the user, if desired. ***Temperature effects and correction:*** As MicrobeMeter is a portable device designed to be placed on a desk at room temperature, or inside a heated incubator, or even to carry around for fieldwork, it is important to identify the stability of its measurements at varying temperatures. This was tested by taking continuous measurements using Hungate tubes containing 10mL of distilled water. Measurements on a 1-minute interval were started and the device was kept at room temperature for 30 minutes, before being moved into an incubator (30°C; Heratherm, Thermo Scientific, MA, USA) for 15 hours and 6 minutes. This data showed approximately 5% decrease when the device was moved from room temperature to 30°C (*Supplementary File 3*). This change was very similar for all ports, indicating that the measurements can be corrected for the temperature effect by using one port with a “blank” sample (i.e., multiplying the measurements of a port using the ratios between the first measurement of that port and the measurements of the “blank” port). This approach was used in the data shown in Figure 3 to 5 and *Supplementary File 3*. ***Temporal stability:*** The data presented in the above section also showed that the temperature corrected measurements of MicrobeMeter are highly stable over time. To test this aspect further, a similar separate test was conducted using pre-heated MicrobeMeter (24 hours at 30°C prior to starting the test) to determine the stability of the measurement over a much longer period of time (65 hours 13 minutes; 3913 measurements; at 30°C). The results showed that the variability among turbidity values was approximately 0.0003 for each port (see *Supplementary File 4* for the data and calculations).

### Biological samples and experiments

The biological results were obtained using different microbes and culturing protocols as follows. ***Cell cultures:*** The following species were used in the presented experiments: *Escherichia coli* K12 (substr. MG1655; originally obtained from the German Culture Collection, DSM18039), *Shewanella oneidensis* MR-1 (originally obtained as a gift from Susan Rosser’s research group at the University of Edinburgh), *Schizosaccharomyces pombe* MBY 102 (genotype: ade6-210 ura4-D18 leu1-32 h+; originally obtained as a gift from Mohan Balasubramanian’s research group at the University of Warwick), *Desulfovibrio vulgaris* (originally obtained from the German Culture Collection, DSM644), and *Methanosarcina barkeri* (originally obtained from the German Culture Collection, DSM800). ***Serial dilutions:*** The *E. coli* and *S. oneidensis* starter cultures were prepared by inoculating 100μL of thawed cryo-stock and liquid culture (grown in lysogeny broth (LB) medium without glucose (18)) into 100mL of LB medium, respectively. These two starter cultures were incubated overnight at 37°C and 30°C, respectively, with 150rpm shaking. Starter culture of *S. pombe* was prepared by picking individual colonies from YEA medium (19) agar plates, dissolving these in 100mL of YEA medium, and incubating for 48 hours at 30°C with 240rpm shaking. After the turbidity values of the starter cultures reached above 2.5, three 33mL-aliquots of each starter culture were transferred into 50mL-centrifuge tubes. The cells were then pelleted by centrifuging at 2000 relative centrifugal force (rcf) for five minutes at 4°C, the supernatants were removed, and each pellet was re-suspended using 23mL PBS. The re-suspended pellets of each organism’s replicate cultures were combined (final volume of 69mL each) and diluted further using PBS to achieve a turbidity measurement between 2.3 to 2.4 before blank subtraction. Turbidity was measured using disposable cuvettes (Fisherbrand, FB55147, Fisher Scientific, UK) and a bench-top spectrophotometer (Spectronic 200, Thermofisher, MA, USA). Two sets of two-fold serial dilutions (1 to 1/16 dilution) were prepared using PBS for each organism. Note that each set contains three technical replicates. The volume of each dilution was 1mL and 10mL in the first and second sets, respectively. The former was used for turbidity measurement using disposable cuvettes, whereas the latter was used for turbidity measurement using Hungate tubes. The turbidity of the well mixed serial dilutions was measured at 600nm using Spectronic 200, Jenway 7305 (Cole-Parmer, Staffordshire, UK) and Novaspec Pro (Biochrom Ltd., Cambridge, UK). Note that the Hungate tube samples were measured only using Spectronic 200 equipped with an adapter for Hungate tubes, and MicrobeMeter. The blanks were prepared using 1mL and 10mL of PBS for the cuvette and Hungate tube samples, respectively. ***Cultures for continuous growth measurements:*** All measurements were done using a pre-heated MicrobeMeter (overnight at 37°C) with three inoculated and one sterile tube (the ‘blank’). After starting the measurements, the MicrobeMeter was moved into a shaking incubator set at 30°C or 37°C (Stuart SI600C, Cole-Parmer, UK or MaxQ 4000, Thermo Scientific, MA, USA). *<u>E. coli experiment</u>*: The starter culture was prepared by inoculating 100μL of *E. coli* K12 from thawed cryo-stock into 50mL of LB medium and incubating at 30°C with 150rpm shaking overnight. Each of the six sterile Hungate tubes containing 10mL of LB medium was then inoculated using 100μL of the starter culture. Two sterile Hungate tubes containing 10mL of fresh LB medium were used for obtaining blank measurements. All tubes were sealed with sterile air-permeable membranes (AeraSeal, LW2783, Alpha Laboratories, UK). The measurements using MicrobeMeter were taken every minute for approximately 30.5 hours at 30°C with 250rpm shaking. Technical replicates (a set of three inoculated tubes and one sterile tube) were also moved into the same incubator and turbidity measurements were taken using Spectronic 200 equipped with an adapter for Hungate tubes every 30 minutes for three hours. *<u>D. vulgaris experiment:</u>* The cell cultures were prepared by inoculating 500μL of a starter *D. vulgaris* culture from its late-log phase. The starter culture was grown anaerobically for four days in a defined medium described in (20), and containing 30mM Na-Lactate and 10mM Na_2_SO_4_. The 500μL inoculum was diluted into sterile Hungate tubes containing 10mL of the same defined medium (three replicates were used, with one un-inoculated blank). Measurements were taken using MicrobeMeter every five minutes for approximately 75 hours at 37°C with 250rpm shaking. *<u>M. barkeri experiment:</u>* The cell cultures were prepared by inoculating 500μL of *M. barkeri* from its late-log phase culture into each of the three sterile Hungate tubes containing 10mL of a defined medium described in (20) (one un-inoculated tube was used as blank). The medium was adapted for *M. barkeri* growth by adding 100μL of 50% (v/v) methanol as sole carbon source and 200μL of 100mM Na_2_S as reducing agent. Furthermore, NaHCO_3_ was omitted as it precipitates during the anaerobic medium heating and degassing processes. Measurements were taken using MicrobeMeter every five minutes for approximately 458 hours at 37°C with 300rpm shaking (high speed shaking was used for avoiding sedimentation of granules).

## ACKNOWLEDGEMENTS

We thank Marco Polin and University of Warwick Physics Workshop for their help with the prototyping of electronic circuit boards, Mohan Balasubramanian for provision of *S. pombe* cultures, Susan Rosser for provision of *S. oneidensis* strain, and Christian Zerfass for preparing *S. oneidensis* starter culture. We also thank past and current members of the Soyer research group for insightful discussions on device development.

## AUTHOR CONTRIBUTIONS

OSS and KS designed the research. OSS contributed to device and electronics design, AVM contributed to device design, JC and TF contributed to culture work, and KS designed the system and electronics, implemented the device, and performed the analyses. OSS and KS wrote the manuscript, which was approved by all co-authors.

## DEVICE AVAILABILITY

The blueprints and additional details (such as data acquisition and analysis software) of MicrobeMeter are made publicly available under a personal and academic non-commercial use licence at www.humanetechnologies.co.uk.

## Supplementary Files (provided as a compressed file)

Supplementary File 1: Data from the technical quality check experiment using pulse-width modulation as explained in the main text.

Supplementary File 2: Data from experiment to determine the response time of MicrobeMeter as described in *Methods* section.

Supplementary File 3: Data from the experiment to identify the effect of temperature on MicrobeMeter measurements as described in *Methods* section.

Supplementary File 4: Data from the experiment to identify the temporal stability of MicrobeMeter as described in *Methods* section.

